# Glycerol is the actuator of integral feedback control in yeast osmotic stress signaling

**DOI:** 10.1101/045682

**Authors:** Suzannah Rutherford

## Abstract

In a 2009 article in Cell van Oudenaarden and colleagues employed elegant experiments and control theory to model perfect adaptation of the yeast osmotic stress response – precise return of turgor pressure to its optimal, steady-state value despite variation in system parameters and the continued presence of osmotic stress. Their data convincingly showed that nuclear signaling and cell volume undergo “robust perfect adaptation” implying that integral feedback must restore their steady state values. However, the authors incorrectly mapped the integrator onto a minimal network that violates assumptions implicit in conventional block diagrams. Using known features of osmotic stress signaling and results presented by the authors, I argue that glycerol concentration – the integral of the rate of glycerol accumulation (synthesis minus leakage) – transforms metabolic energy into an increased osmolarity that drives water influx and restoration of turgor pressure. I show how integral feedback control actuated through glycerol synthesis is logically positioned to provide perfect adaptation and robustness in hyperosmotic stress responses.

## Introduction

Robust perfect adaptation is a property of biological feedback control systems that precisely maintains steady-state homeostasis of vital functions in the continued presence of disturbances such as environmental stress or, in the case of desensitization, increases the dynamic range of detection for external signals such as pheromones or environmental nutrients (Yi et al., 2000). In their article “A systems level analysis of perfect adaptation in yeast osmoregulation” Muzzey et al. beautifully demonstrated perfect adaptation of nuclear signaling, cell volume, and thus membrane turgor pressure in osmotic stress (Muzzey et al., 2009). Control theory shows that perfect adaptation predicts and is predicted by integral feedback control (Yi et al., 2000). Muzzey and colleagues argue convincingly and with clever experiments that their data support a single integrating mechanism (or integrators acting in parallel, but not in series). The authors use this point to identify on a minimal network model the location of the integral feedback control mechanism(s). I argue here that the model was incorrect and that a major conclusion of the paper, the inferred location of the integrator, should be revisited.

Analyses of complex control systems depend critically on an accurate working model with clearly defined topology (Riggs, 1970; Romagnoli and Palazoglu, 2006).

Here I show that the minimal representation of the osmotic stress response presented in Muzzey et al. violates assumptions implicit in conventional block diagrams (page 115 (Riggs, 1970)). Using known features of osmotic stress signaling and results presented by the authors, I argue that integration of the summed rates of glycerol synthesis and leakage (amount per time) into an accumulating intracellular concentration of glycerol (amount per volume) provides integral feedback control, perfect adaptation, and robustness to the hyperosmotic stress response. In contrast with localization of the integrator in the HOG (high osmolarity glycerol) dependent branch of the pathway as proposed by the authors (Muzzey et al., 2009), I argue instead that glycerol concentration is the integrator. By responding to both HOG dependent and HOG independent processes intracellular glycerol concentration integrates the cell’s response to increased external osmolarity; by directly driving water influx, osmolarity, and restoration of turgor pressure intracellular glycerol concentration actuates the response.

## Results

Figure 1 shows a detailed block diagram representation of feedback control based on features of the osmotic stress response using as a general template the feedback control system presented in Figure 10.4 of Romagnoli and Palazoglu (Romagnoli and Palazoglu, 2006). In this representation each block depicts a complete, unidirectional subsystem whose output variables are uniquely and completely determined by their input variables (Riggs, 1970).

It is well documented that the primary survival function of the osmotic stress response is to restore turgor pressure through increased synthesis and accumulation of glycerol (reviewed in (Hohmann et al., 2007)). As in the minimal representation presented by Muzzey et al. I start with measurement of turgor pressure, the controlled variable (y). By contrast with the minimal representation, in the detailed model turgor pressure is transduced into biologically meaningful signals through multiple known (and possibly additional unknown) measuring functions (g_m_^i^). These are upstream signal transduction proteins with effects on turgor pressure adaptation (Hohmann, 2002; Hohmann et al., 2007; Rep et al., 1999). For example, 1) the osmosensitive glycerol channel Fps1 closes in response to decreased turgor pressure, 2) activating conformational changes in the osmosensor proteins Sho1 and Sln1 respond to decreased turgor pressure to initiate the high osmolarity glycerol (HOG) pathway and 3) inactivating conformational changes in other proteins may respond to directly or indirectly to reduced turgor pressure or downstream changes (e.g. cytoplasmic crowding (Miermont et al., 2013)).

For simplicity, I assume that the steady-state activities of the sensor proteins (plausibly their relaxed conformations) are zero when turgor pressure is at its optimal value, however non-zero steady-state deviations are accommodated by the general model (Romagnoli and Palazoglu, 2006). If steady-state activities are zero the cellular measurements of turgor pressure (y_m_) are equivalent to their deviations from steady-state (errors; e^i^) for all sensors. Following standard convention, the e^i^ are fed into the controller (grey block) through the detailed input-output functions shown. The output of the glycerol synthetic machinery in the controller is the controlled variable (c), the rate of glycerol accumulation. The rate of glycerol accumulation is the summation of the rate of synthesis, provided through regulation of the pathway at several levels, and the opposing rate of glycerol leakage, through the Fps1 channel (Remize et al., 2001). The intracellular concentration of glycerol is the manipulated variable (m).

Mapping the minimal model of Muzzey et al. (Figure 2A) onto the detailed representation in Figure 1 shows 1) how their subsystems H and I violate single-input single-output (SISO)assumptions and are therefore invalid transfer functions, 2) where the subsystems overlap, and 3) how their minimal model is not isomorphic with the detailed model of the osmotic stress response including known features discussed in their article (Figure 2B; a minimal model consistent with the detailed block diagram of the response is given in Figure 2C). Moreover, simply by definition glycerol concentration is an integrator: the cumulative amount (glycerol per cell volume) is proportional to the integral of the rate of glycerol accumulation (amount per time). Glycerol is downstream location of the integrator suggested by Muzzey et al., who demonstrated that the osmotic stress response network in yeast must contain exactly one integrator acting in series (or possibly more than one integrator acting in parallel). Therefore the locus of integration must reside in the intracellular glycerol concentration itself, and not the HOG-dependent block of the network as concluded by the authors (Muzzey et al., 2009).

At steady-state the intracellular concentration of glycerol is by definition constant. If the concentration of glycerol is constant, then net glycerol accumulation must be zero, and the rate of glycerol leakage must be equal to its rate of synthesis. As shown by Muzzey et al., similar logic applies to all upstream components; if the integrator is the most downstream element in the network (relative to the input of a disturbance in external osmolarity), all error deviations, Hog1 nuclear enrichment, and steady-state viability must also display perfect adaptation (Muzzey et al., 2009). Indeed, we observed perfect adaptation of viability before and after adaptation to an osmotic challenge (correlation between early mortality and recovery of viability of 50 different yeast strains is over 0.98; http://biorxiv.org/content/early/2016/03/07/039982). Given enough time (depending on the time constants of each response) cells adapt to the hyperosmolar media with increased intracellular glycerol concentrations. Once adaptation occurs and turgor pressure is restored, sensors relax and all deviations from steady state activity return to zero.

## Discussion

The molecular mechanisms behind error sensing have been a source of mystery to bioengineers (e.g. “the molecular mechanisms behind error sensing are little understood” (Xiao et al., 2009)). However, it is easy to imagine how changes in turgor pressure could be translated into conformational shifts between relaxed and activated states of proteins as an error sensing mechanisms in the osmotic stress response (Rutherford and Zuker, 1994). Conformational distortions caused by less than optimal turgor pressure are thought to activate the high osmolarity glycerol (HOG) pathway in proportion to osmotic stress. Similar activation or loss of activity of cytosolic proteins plausibly occur in conditions of molecular crowding (Miermont et al., 2013). In that case, restoration of cell volume and turgor pressure would allow proteins to return to pre-stress steady conformations and levels of activity. Finally, the conclusions of this analysis are generalizable. The intracellular concentration of any biomolecule is the integral of its positive rate of accumulation, and provides a possible source of integral feedback.

These are also plausible actuators of homeostasis, converting the energy of cellular metabolism into the energy inherent in concentration gradients for other concentration dependent processes.

In addition to revealing potentially general features of integral feedback control in biology, the block diagrams in Figures 1 and 2c show how the osmotic stress response may work over different time scales and levels of stress. Upon a shift to hyperosmotic media, the initial response is the rapid closure of the constitutive leakage channel Fps1, closing the shortest negative feedback loop in Figure 1 and effectively removing the negative input to the controller in the circuit shown in Figure 2c. Figure 1 also shows clearly and intuitively how a successive activation of less sensitive sensors with longer time scales and additional sources of negative feedback on glycerol accumulation could occur, consistent with the observed longer delays and increasingly stronger adaptive responses proportional to the degree of osmotic stress (Hohmann et al., 2007).

## Figure Legends

**Figure 1. The yeast osmotic stress response as an error actuated linear control system.** At steady-state (time 0^-^) turgor pressure (y) is at its pre-stress value and osmostress sensitive proteins (sensors 1-5) are in their relaxed, non-induced conformations, with all error deviations (e) equal to 0. An abrupt step in external osmolarity (d) transiently alters turgor pressure, the controlled variable (y). The change in turgor pressure (de)activates sensor proteins that transduce the signal to downstream components. The canonical osmotic stress response pathway is the high osmolarity glycerol (HOG) MAP kinase cascade (H; purple) activated by more-sensitive and less-sensitive sensors Sln-1 and Sho-1 (Hohmann et al., 2007). Dual phosphorylation of the downstream MAP kinase Hog1 in the cytoplasm activates the glycerol synthetic pathway (G; orange). Later, active Hog-1 is translocated to the nucleus (D; blue; nuclear Hog1 dependent functions), where it controls transcription and synthesis of GPD-1, the rate limiting enzyme in glycerol synthesis(Remize et al., 2001). More rapid, nuclear Hog-1 independent activities (I; green) also increase glycerol synthesis and retention. Upon activation by osmotic stress the Fps-1 glycerol channel, normally open, immediately closes. The osmosensitive kinase Ypk1 responds to osmotic stress by increasing the activity of GPD1 (Lee et al., 2012), and the function of proteins that are sensitive to osmotic stress and/or cytoplasmic crowding is impaired. The loss of activity of proteins that serve essential functions plausibly cause reduced fitness and loss of viability, activating general stress responses including the general stress response transcription factor Msn-2 whose target is (among others) GPD1 (Gasch et al., 2000). Processes initiated at sensors 1-5 are activated by successively higher levels of osmotic stress and act on successively longer timescales. For example, less severe osmotic stress has a very short response time and may not activate Sho-1 (e4) or damage cellular proteins (e5). Linear relationships between Laplace transformed of inputs and outputs of the transfer functions are assumed (Riggs, 1970; Romagnoli and Palazoglu, 2006).

**Figure 2. Minimal model violates single-input single-output (SISO) assumptions of conventional block diagrams. A.** The minimal circuit model of Muzzey et. al. with 4 blocks identified as 1) the H subsystem that contains the MAP kinase cascade and “links an osmotic disturbance at the membrane with Hog1 nuclear enrichment”, 2) the D subsystem that contains “Hog1 dependent mechanisms that promote glycerol accumulation including the transcriptional activation of genes encoding glycerol producing enzymes and interactions initiated by Hog1 in the cytoplasm or nucleus that lead to increased glycerol synthesis”, 3) the I subsystem that contains “Hog-independent mechanisms that contribute to osmolyte production” and 4) the G subsystem representing “metabolic reactions involved in glycerol synthesis and any other reactions that contribute to glycerol accumulation” (Muzzey et al., 2009). A mathematical implementation of the model shows how turgor pressure can return to pre-stimulus values even in the continued presence of osmotic stress (Muzzey et al., 2009) but does not prove that the model correctly reflects the biology. **B.** A revised block diagram showing in grey additional links indicated in the text of Muzzey et al. but not shown in their model that violate SISO assumptions. For example, subsystem H has one input (turgor pressure) but 2 outputs (activated Hog1 in the nucleus which increases transcription of GPD1 (blue in Figure 1) and activated cytoplasmic Hog1, which is believed to act through Pfk2c in combination with other outputs to increase glycerol pathway activity (orange in Figure 1)). (Indeed the revised circuit must include 2 additional summation points not in the original model.) Subsystem I, nuclear Hog1-independent functions has three independent outputs: 1) the general stress response inducing Msn2/4, which further activates GPD1 transcription (Boy-Marcotte et al., 1998; Gasch et al., 2000)(ref), 2) the Fps1 leakage channel closing counteracts glycerol synthesis(Hohmann, 2002), and 3) increased Gpd1 activity through nuclear Hog1-independent mechanisms is summed with Hog1-dependent increases in Gpd1 synthesis to promote glycerol pathway activation (e.g. Ypk1 (Lee et al., 2012), Pfk26 (Dihazi et al., 2004) and as reviewed by (Hohmann, 2002; Hohmann et al., 2007; Saito and Posas, 2012)). **C.** Minimal model that is topologically equivalent to the model in Figure 1. Model includes a single controller (grey) with input turgor pressure and output glycerol accumulation, a single integrating mechanisms converting summed rates of glycerol synthesis and loss to intracellular glycerol concentrations and two (groups of) sensors. The positive (sensors 2–5) and negative (sensor 1) mechanisms promote glycerol synthesis or leakage. For comparison, the locations of the four subsystems depicted in Figure 2A are shown.

